# Ecological advantages and evolutionary limitations of aggregative multicellular development

**DOI:** 10.1101/255307

**Authors:** Jennifer T. Pentz, Pedro Márquez-Zacarías, Peter J. Yunker, Eric Libby, William C. Ratcliff

## Abstract

All multicellular organisms develop through one of two basic routes: they either aggregate from free-living cells, creating potentially-chimeric multicellular collectives, or they develop clonally via mother-daughter cellular adhesion. While evolutionary theory makes clear predictions about trade-offs between these developmental modes, these have never been experimentally tested in otherwise genetically-identical organisms. We engineered unicellular baker’s yeast (*Saccharomyces cerevisiae*) to develop either clonally (‘snowflake’, Δ*ace2*), or aggregatively (‘floc’, *GAL1*p::*FLO1*), and examined their fitness in a fluctuating environment characterized by periods of growth and selection for rapid sedimentation. When cultured independently, aggregation was far superior to clonal development, providing a 35% advantage during growth, and a 2.5-fold advantage during settling selection. Yet when competed directly, clonally-developing snowflake yeast rapidly displaced aggregative floc. This was due to unexpected social exploitation: snowflake yeast, which do not produce adhesive FLO1, nonetheless become incorporated into flocs at a higher frequency than floc cells themselves. Populations of chimeric clusters settle much faster than floc alone, providing snowflake yeast with a fitness advantage during competition. Mathematical modeling suggests that such developmental cheating may be difficult to circumvent; hypothetical ‘choosy floc’ that avoid exploitation by maintaining clonality pay an ecological cost when rare, often leading to their extinction. Our results highlight the conflict at the heart of aggregative development: non-specific cellular binding provides a strong ecological advantage – the ability to quickly form groups – but this very feature leads to its exploitation.

## Introduction

The evolution of complex life on Earth has occurred through key steps in which formerly autonomous organisms evolve to become integral parts of a larger, higher-level organism^1–5^. These have been termed Major Transitions in Evolution^5^, or Evolutionary Transitions in Individuality^2,6^, one example of which is the transition from uni- to multi-cellularity. Multicellularity has evolved in at least 25 times in organisms as diverse as bacteria^7,8^, archaea^9^, and among deeply-divergent lineages of eukaryotes^10,11^.

There are two basic modes of multicellular development. Cells can ‘stay together’ after mitotic division, resulting in clonal development if the life cycle includes a genetic bottleneck^7,12^. Alternatively, potentially unrelated cells can ‘come together’ via aggregation, which occurs in a few groups of terrestrial microorganisms^13,14^. Clonal development is thought to possess several advantages over aggregation for multicellular construction. First, under clonal development, cells comprising the multicellular organism have a high degree of genetic relatedness^15^, which aligns the fitness interests of individual cells, facilitating the evolution of cooperative traits (*e.g.*, division of labor). Additionally, clonal development limits the potential for evolutionary conflict, as there is little standing genetic variation within an organism for selection to act on^16–19^. Through the same mechanism, clonal development stifles opportunities for the evolution of parasitic cell lineages that infiltrate and exploit functional organisms^20^. Second, organismal clonality facilitates cluster-level selection. Genetic uniformity among the cells in a group results in a direct correspondence between emergent multicellular traits and heritable information (primarily genes) responsible for generating these traits^21,22^. Variation in the identity and frequency of different genotypes of cells within aggregates across multicellular generations undermines the heritability of emergent multicellular traits. Further, clonal development facilitates the shift from selection acting among cells to whole groups, simultaneously minimizing within-group genetic variation (thus largely preventing within-group selection) and maximizing between-group genetic variation^16^. Perhaps because of these benefits, the majority of independently-evolved multicellular lineages develop clonally.

Yet aggregative development possesses a unique (but largely unappreciated) advantage: multicellular bodies can form far more rapidly^12^. If a group is formed via the ‘staying together’ of cells after division, then its formation occurs by growth, causing the time required for body formation to scale with cellular generation time and organism size. In contrast, aggregation can occur far more rapidly. For example, aggregation of *Dictylostelium* into a multicellular mound can occur just 4-6 hours after starvation^23^, and flocculation of yeast can occur in seconds^24^. Indeed, aggregative development is common in organisms that rapidly switch from unicellular to multicellular life history strategies upon sudden environmental change (*e.g.*, starvation in *Dictylostelium discoideum*^25^ and *Myxococcus xanthus*^26^). Aggregation may also bring together cells with complementary properties, taking advantage of mutualistic interactions^27–29^, but the evolutionary stability of this interaction generally requires a mechanism to limit social exploitation, such as host sanctions^30,31^ or partner fidelity across generations^32^.

The origin of complex life cannot be understood in the absence of evolutionary mechanism. It thus is imperative that we understand how basic mechanisms of multicellular development effect the subsequent evolution of multicellular complexity. Mathematical modeling^12,19,21,22,33–36^ and experiments in diverse systems^20,37–41^ have generated consistent and robust predictions for the evolutionary consequences of variation in developmental mode. Yet because no model organisms develop through both routes, no experiments have directly compared ecological vs. evolutionary trade-offs between aggregative and clonal development. Here, we circumvent this historical limitation by engineering unicellular yeast (*Saccharomyces cerevisiae*) so that they form multicellular groups via either clonal development or aggregation.

The yeast *S. cerevisiae* can aggregate to form large clumps consisting of thousands of cells termed ‘flocs’. Aggregation occurs via a lectin-like bonding between cell surface FLO proteins and cell wall sugars in adjacent cells^38,42^. Flocs preferentially form among mutual FLO^+^ cells; FLO^−^ cells tend to be excluded from the group^43^. However, genetically-diverse strains can join a floc if they are FLO^+^ (Figure 1a). In contrast, ‘snowflake yeast’ develop clonally, forming multicellular groups as a consequence of failed septum degradation after cytokinesis^44^; Figure 1A). When a cell-cell connection is severed, the group produces a viable propagule. This propagule experiences a single-cell genetic (but not physiological) bottleneck, as the most basal cell in the propagule is the mitotic parent of every cell in the group^44^.

**Figure 1.**
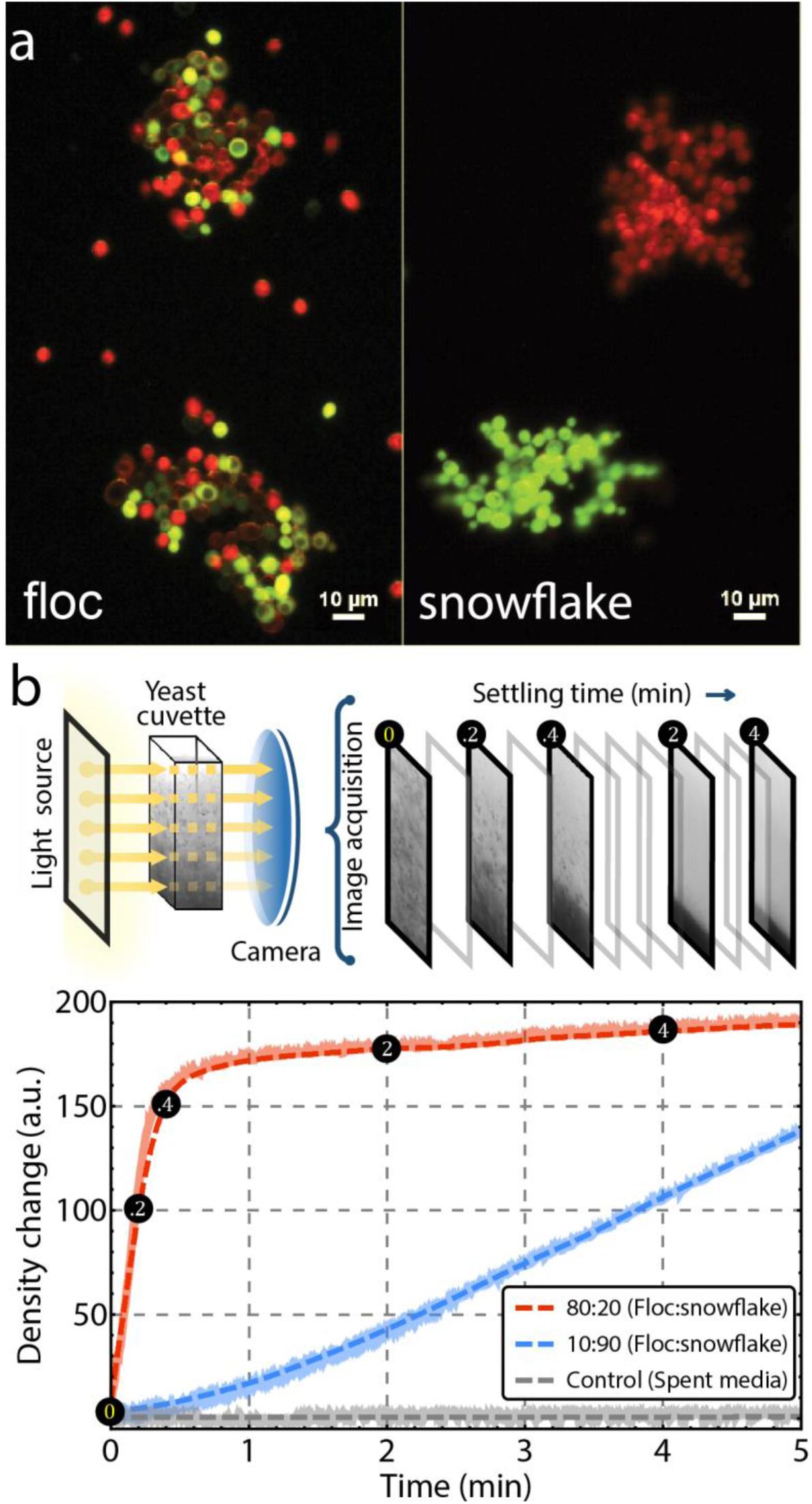
Synthetic yeast system to study clonal and aggregative multicellular development. a) Synthetically created floc and snowflake yeast (*FLO1* insert and *ace2* knockout, respectively) labeled with either a red or green fluorescent marker. Both strains were created from the same unicellular ancestor. Flocs may be genetically-diverse, while snowflake yeast form clonal clusters. b) Settling rate was measured using high-resolution video acquisition of back-illuminated yeast cultures over 5 minutes of settling. Individual pixel intensities, which correlate to yeast density, were used to measure the rate of density change (see Supplementary Movie 1). Raw density data (shadowed lines) was smoothed with a Savitzky-Golay smoothing function (dashed line) and the maximum slope of these dynamics is calculated as the settling rate. Shown are the density dynamics of fast (80:20 Floc:Snowflake) and slow (10:90 Floc:Snowflake) co-cultures, as well as a cell-free control where no density change is expected.

Engineered isogenic floc and snowflake yeast were constructed from a common unicellular ancestor. They were grown in a fluctuating environment, 24 hours of shaking incubation followed by selection for rapid sedimentation, that favors a rapid transition from unicellularity (providing the highest growth rates) to multicellularity (increasing survival during settling selection). Aggregation was a superior strategy in monocultures: floc yeast, which spend most of the growth phase as unicells or small groups, grew 35% faster than snowflake yeast, but rapidly formed large flocs during settling selection, settling 2.5 times as fast as snowflake yeast. Yet in competition, snowflake yeast rapidly outcompete floc, the result of an unexpected social interaction. Despite being FLO^−^, snowflake yeast embed themselves within floc clusters, making up a disproportionately high fraction of the biomass within flocs. Spatial analysis of chimeric aggregates demonstrates that snowflake yeast are uniformly, not randomly, distributed within the floc, suggesting a simple physical interaction between floc and snowflake is necessary for the formation of chimeric aggregate clusters. In principle, this parasitism could be prevented if floc evolved a partner choice mechanism, excluding heterospecific genotypes. We examined the invasion of such a ‘choosy’ floc genotype using mathematical modeling. In our model, selective binding is ecologically costly, as there is an advantage for individual cells to form groups with as many other cells as possible (this way they form the largest groups). Rare choosy floc is therefore unable to invade permissive floc, snowflake yeast, or a population consisting of both. Because choosy floc’s aggregative performance is strongly frequency dependent, it should perform poorly (relative to a permissive floc) in genetically-diverse populations. This ecological cost may limit the evolution of strong kin recognition during aggregative development, paving the way for persistent evolutionary conflict.

## Results

There are two important life history traits that affect fitness in our fluctuating environment: growth during 24 h batch culture and settling rate during settling selection^45–47^. To measure settling rate, we developed a novel method to quantify the dynamical effects of aggregation and settling in real time (Figure 1b, Supplementary Movie 1; see methods section for details). Floc yeast are superior in both traits. First, floc yeast settle 2.5 times as fast as snowflake yeast, rapidly forming large aggregates during settling selection (Figure 2a; Movie S2; *t_8_* = 9.82, *p*<0.0001, two-tailed t-test). In direct competition, floc yeast outcompete snowflake yeast over one 24 h growth cycle (Figure 2b). This is likely a consequence of nutrient and oxygen limitation in snowflake clusters, which, in contrast to floc yeast, are always multicellular.

**Figure 2.**
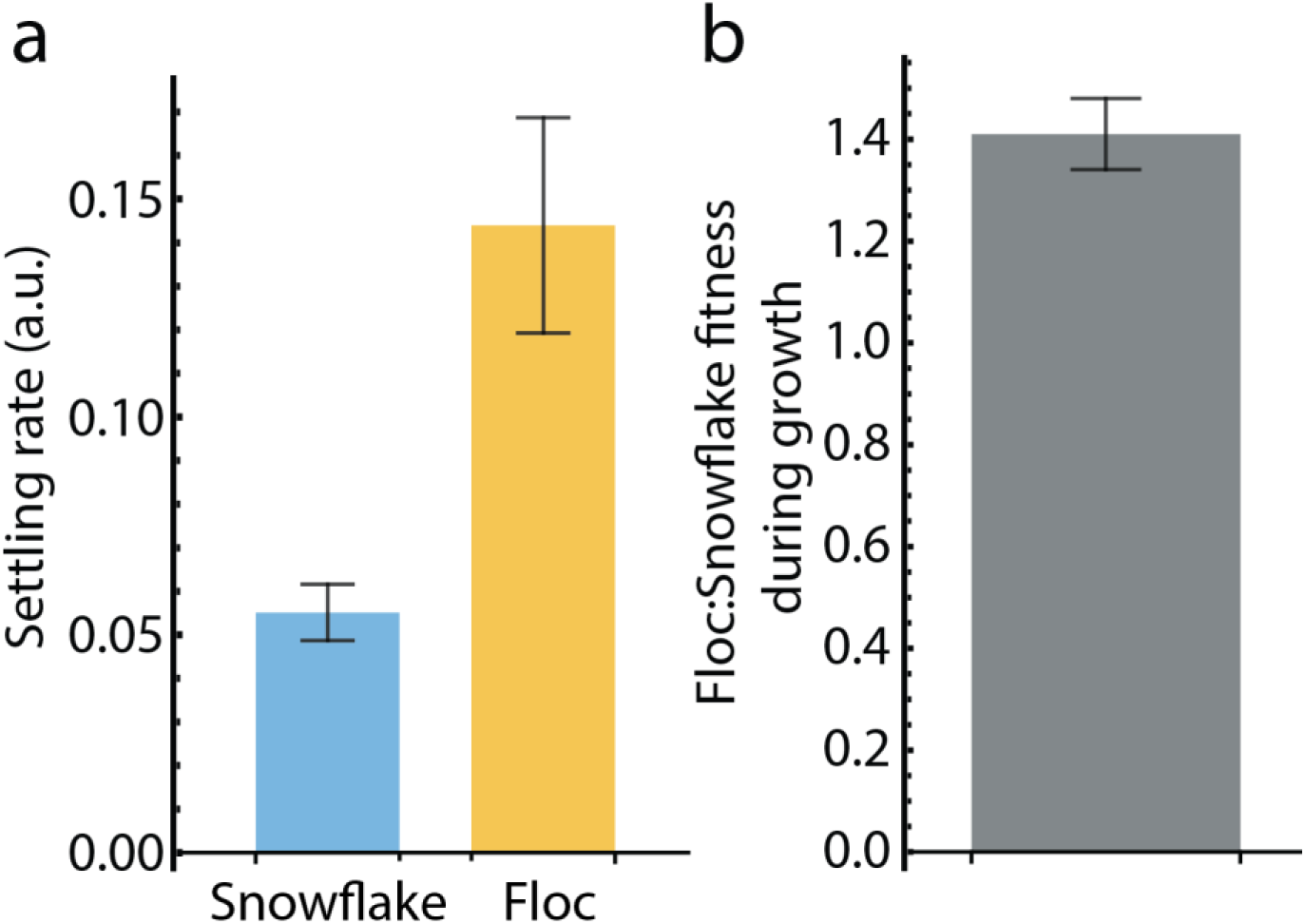
Aggregative floc yeast are more fit than clonally-developing snowflake yeast in an environment favoring rapid group formation. Floc yeast are superior in two important life history traits that affect fitness in our experimental system. a) Floc yeast settle 2.5 times faster than snowflake yeast (*t_8_* = 9.82, *p*<0.0001, two-tailed t-test). Error bars are standard deviation (n=8). b) Floc yeast outcompete snowflake yeast over one 24 h growth period. Fitness was measured as the ratio of Malthusian growth parameters^70^ for one 24 h period. Error bar is standard deviation (n=5).

Co-culturing floc and snowflake yeast introduced markedly different behaviors. The settling rates of mixed populations increased dramatically (Figure 3a), and was highest when snowflake yeast were at an intermediate frequency (20-50%; F_10,33_ = 25.5; *p* < 0.0001; ANOVA, pairwise differences assessed with Tukey’s HSD with α = 0.05). To examine the effects of co-culture on fitness, we performed a series of competition experiments (two rounds of growth and settling) across a range of starting snowflake frequencies, from 1% to 99%. Surprisingly, snowflake yeast were more fit than floc in all competitions, and their fitness was highly frequency-dependent. When snowflake yeast were rare (starting at 1% of the initial culture biomass), they had a small competitive advantage over floc (Figure 3b). This increased dramatically when they were more common (10-20% of initial starting biomass), declining until snowflake yeast reached 80%. Flocculation was impeded when snowflake yeast constituted >80% of the population, allowing multicellular snowflake yeast to compete against largely unicellular floc, causing their relative fitness to again increase dramatically (Figure 3b&d). These dynamics appear to be the result of an unexpected interaction: when mixed together, snowflake yeast and floc form chimeric clusters during the settling phase of the experiment (Figure 3c).

**Figure 3.**
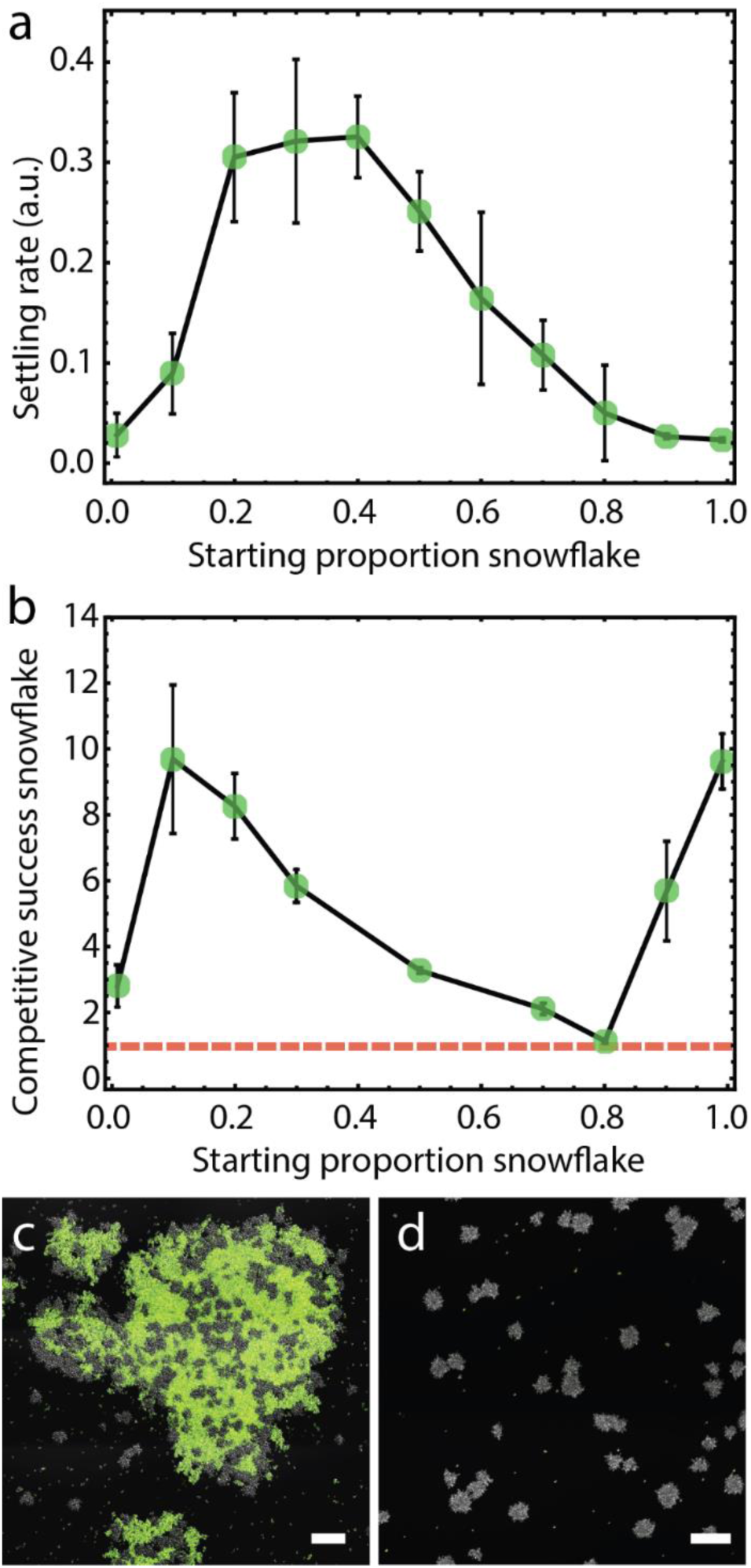
Co-culturing floc and snowflake yeast. a) Mixed populations settle more rapidly than snowflake yeast or floc alone. Settling occurs the most rapidly at intermediate frequencies (20-50%; F_10,33_ = 25.5; *P* < 0.0001; ANOVA followed by Tukey’s HSD). Error bars are standard deviation of four biological replicates, settling rate units are arbitrary. b) We measured the competitive success of snowflake yeast across two rounds of growth and settling. Snowflake yeast were more fit than floc at all genotype frequencies. Error bars are standard deviation of five biological replicates. (c). Snowflake yeast form chimeric aggregates with floc. Shown are snowflake yeast and GFP-tagged floc yeast starting at an initial inoculation ratio of 30:70 snowflake:floc-GFP (c) or 99:1 (d). Note that floc are below the concentration threshold required for aggregation, existing as unicells. Scale bars are 100 µm.

To determine which phase of the periodic environmental regime (*i.e.*, growing vs. settling) favored snowflake yeast during competition with floc, we measured snowflake yeast competitive success across one culture cycle. Consistent with earlier experiments (Figure 2c), snowflake yeast lost to floc over one 24 h growth cycle (Figure 4). Snowflake yeast fitness during growth was negative frequency dependent (y = −0.005x + 0.91, *p*<0.0001, linear regression). This is likely a consequence of overall nutrient consumption rates. When slower-growing snowflake yeast make up a larger fraction of the population, they consume resources less quickly, extending the time over which their floc competitors can compound their growth rate advantage. In contrast to growth, however, snowflake yeast possessed an advantage during settling selection (Figure 4).

**Figure 4.**
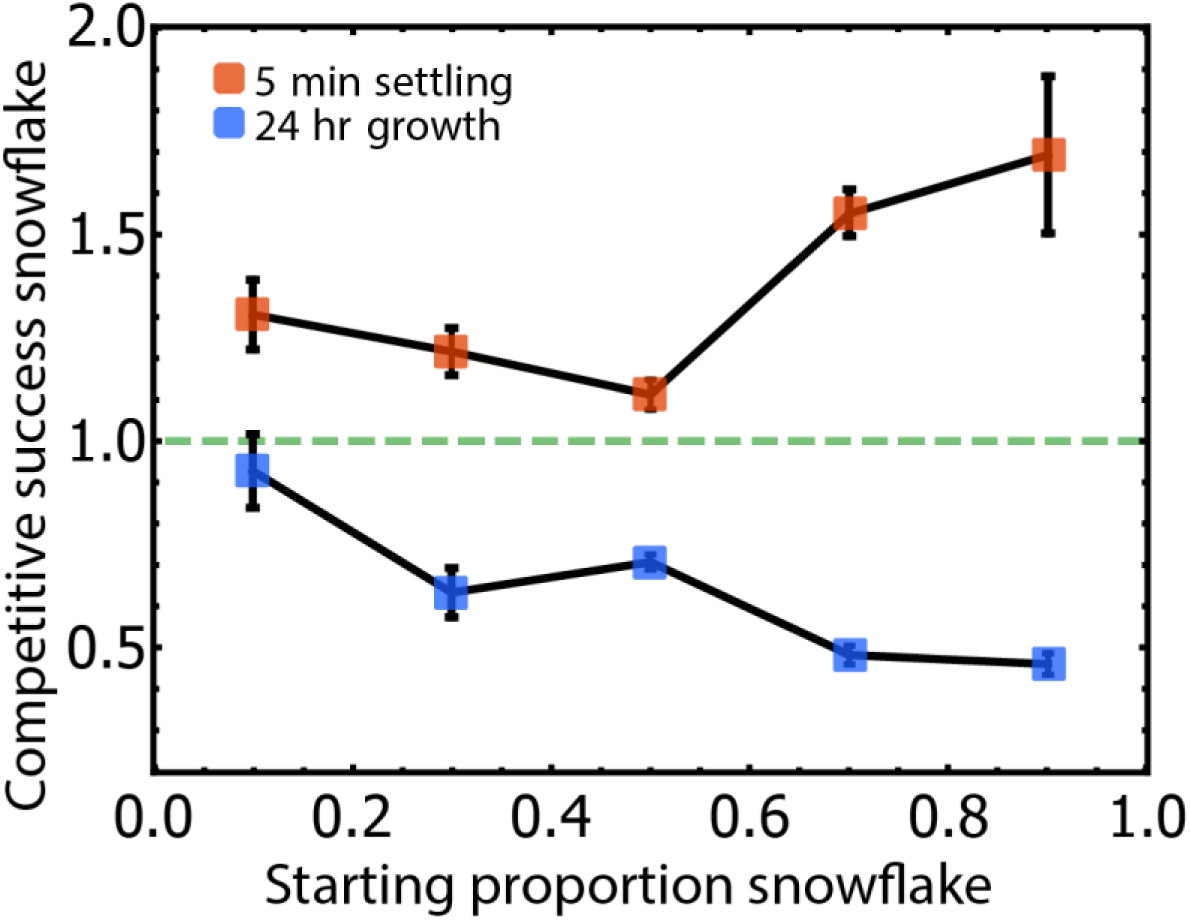
Snowflake yeast outcompete floc during settling selection when forming chimeric aggregates. We examined the competitive success of snowflake yeast in competition with floc during both growth (over 24 h of culture) and settling selection (5 minutes at 1 g). Snowflake yeast had lower fitness at all starting genotype frequencies during the growth phase of the culture, yet had higher fitness during settling selection. This is in stark contrast to what we observe in pure culture, where floc yeast settle 2.5 times as quickly as snowflakes (Figure 1b). Error bars are standard deviation of five biological replicates.

One way that snowflake yeast could gain an advantage during settling selection is if they are over-represented in large, fast-settling chimeric aggregates. This would be unexpected, as *FLO1* yeast preferentially adhere to other floc cells, efficiently excluding non-flocculating unicells from flocs^43^ (Supplementary Figure 1). We imaged co-cultures in which snowflake yeast were either rare (20% initial biomass) or common (80% initial biomass). Surprisingly, snowflake yeast were overrepresented in chimeric flocs (*i.e.*, groups larger than the largest individual snowflakes; Figure 5c) at both genotype frequencies.

**Figure 5.**
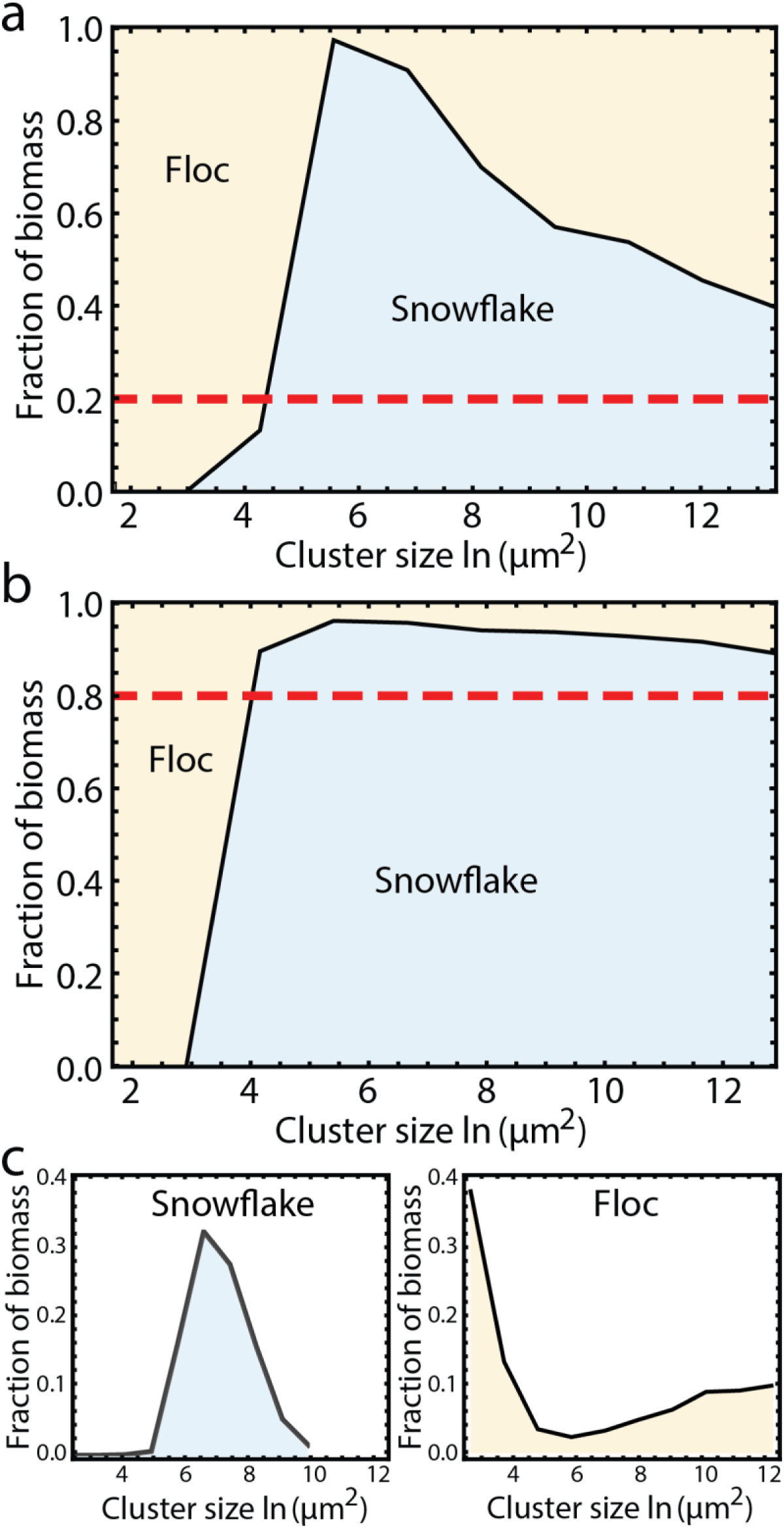
Snowflake yeast are overrepresented in large chimeric aggregates. Snowflake yeast constitute a larger fraction of the biomass within large flocs than is expected by their overall population frequency (dashed line). Shown are snowflake yeast at 20% (a) and 80% (b) overall frequency. c) Size distributions of pure snowflake and floc cultures.

One feature of chimeric aggregates that stands out is the appearance of a relatively uniform distribution of snowflake yeast within the aggregate (Supplementary Figure 2). We rarely see large patches of pure floc cells, and never see large patches of just snowflake yeast. To quantify the spatial distribution of snowflake yeast within chimeric aggregates, we first measured the spatial autocorrelation function (Moran’s *I*). We found that the correlation length is similar in size to the cluster radius (14.1 +/− 0.2 µm, 14.2 +/− 0.2 µm, 13.9+/− 0.1 µm, 11.7+/−1.3 µm for 30%, 20%, 10%, and 1% snowflake yeast, respectively). We next measured the pair correlation function, *g(r)*, which measures the probability of finding two clusters separated by a given distance (Supplementary Figure 3), normalized by a random distribution at the same density. We find that the distribution of snowflake yeast clusters is highly structured within aggregates. Clusters are unlikely to be found very close to each other; specifically, clusters are less likely to be found with a center-to-center separation less than or equal to 1.3 times their diameter than expected by random chance. Relatedly, clusters are more likely to be found with center-to-center separations between 1.3-1.9 times their diameter than one expected by random chance. Thus, the distribution of clusters within an aggregate is more evenly dispersed than would be expected by a random mixing of genotypes. This even dispersal suggests that snowflake yeast are capable of binding to a patch of floc cells, but not a patch of snowflake yeast, during aggregate formation. Consistent with this hypothesis, floc appear to act as an adhesive, binding together snowflakes (Supplementary Figure 4). We do not see any evidence of direct snowflake-snowflake adhesion. This analysis shows that snowflake yeast join chimeric aggregates more efficiently than floc yeast, despite the fact that floc yeast can stick to both floc and snowflake yeast, while snowflake yeast can only stick to floc yeast.

A classic solution to social conflict in aggregating multicellular organisms is kin recognition, allowing individuals to prevent cheating by only joining groups with close relatives^48–51^. Here, we examine whether a self-recognition mechanism would help flocculating yeast outcompete snowflake yeast by constructing a mathematical model (see Methods). Briefly, we assume that there are three types of yeast: a snowflake yeast strain (*S*), a “choosy floc” (*C*) that uses a self-recognition mechanism to adhere just to clonemates, and a “permissive floc” (*P*) that has no such self-recognition mechanism, adhering to both permissive floc and snowflake yeast. We simplify our analysis by focusing strictly on the role of self-recognition in the formation of groups. Thus, we assume that after some initial period of population growth, there is an aggregation phase in which cells stop reproducing and the flocculating yeast aggregate to form groups. Rather than modeling the complex dynamics of group size and shape during settling selection, we make the simplifying assumption that only the largest groups survive. While floc yeast rapidly form groups, increasing in size as a function of time (Supplementary Figure 5), snowflakes themselves do not change in size (as there is no growth; Supplementary Figure 5), though they may join aggregates with permissive floc. When floc are growing at higher density, it takes less time to form groups that can outcompete snowflake yeast during settling selection (Supplementary Figure 5).

We consider all pair-wise competitions between permissive floc, choosy floc, and snowflake yeast for different starting genotype frequencies (Figure 6a-c). For each competition, we simulate the aggregation process and then select 10% of the population from the largest groups (selection that is roughly analogous to the experimental protocol). Snowflake yeast are overrepresented within large, fast-settling flocs (recapitulating our experimental data; Figure 6), allowing them to outcompete permissive floc regardless of their starting frequency. We also find that the largest chimeric aggregates, representing the fastest-settling aggregates, form with intermediate frequencies of snowflake yeast (peaking at 40% S; Supplementary Figure 7). This is similar to our experimental data (Figure 3a), where the fastest-settling aggregates are also found at intermediate frequencies (20%-50%). In contrast, if snowflake and choosy floc compete, then choosy floc increases in abundance whenever it is more than ~60% of the population (though the precise frequency depends on model parameters, like density, aggregation time, and binding probability; Supplementary Figures 6, 8, 9). Thus, neither snowflake yeast nor choosy floc can invade each other when rare. Finally, since permissive and choosy floc behave the same in the absence of snowflake (they do not co-aggregate), their dynamics are entirely frequency dependent and neither can invade from rare.

**Figure 6.**
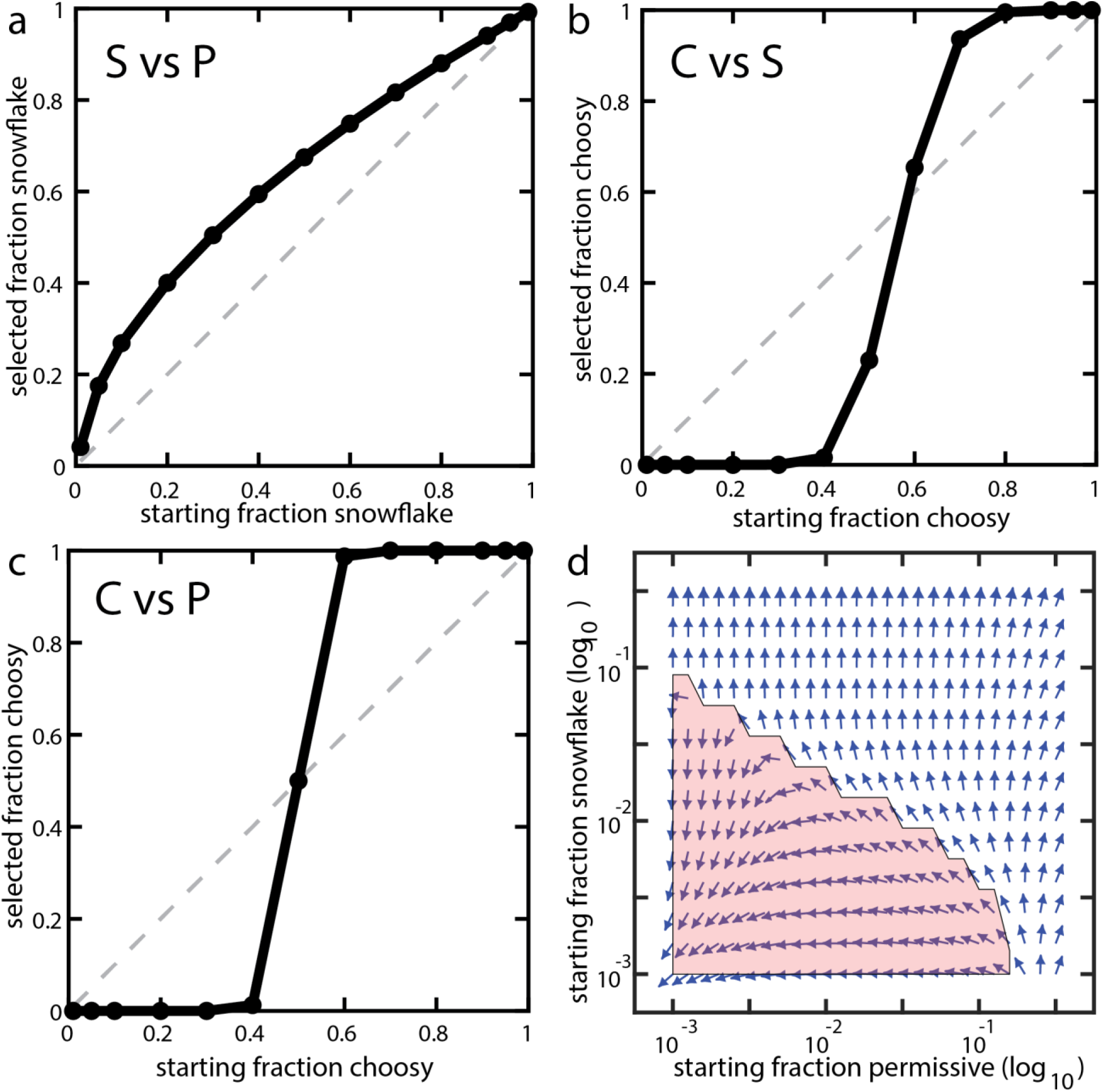
Modeling the dynamics of kin recognition in floc yeast. a) Snowflake yeast, *S*, were capable of displacing permissive floc, *P*, at all frequencies during settling selection. In contrast, the fitness of choosy floc, *C*, (b) and permissive floc (c) were both strongly positively frequency dependent. d) Phase portrait showing the changes in *P* and *S* after one round of settling selection in competition with *C*. Arrows show the direction of change in proportion of *S* and *P* as a function of different starting frequencies. When *P* and *S* start out above a critical threshold, they displace *C*; otherwise, *C* displaces them (red highlight).

In a three-way competition, snowflakes can invade populations of choosy floc with the help of permissive floc (Figure 6d; see results from longer durations of aggregation in Supplementary Figure 8). By forming large, fast-settling chimeric aggregates, mixtures of snowflake and permissive floc can outcompete choosy floc (Figures 6d; Supplementary Figure 8). Of course, this is an unstable alliance, as the exploitation of permissive floc will ultimately result in a monotypic population of snowflake yeast (Figures 6a&d; Supplementary Figure 8). Sometimes, however, this social exploitation of floc is costly for snowflake yeast. When snowflake and permissive floc are below the threshold required to displace choosy floc, exploitation of permissive floc results in a rapid deterioration of their ability to make large chimeric aggregates, to the detriment of both snowflake and permissive floc (Figure 6d; Supplementary Figure 8).

A simple extrapolation of our model highlights the cost of kin discrimination during aggregative development. Consider a genetically-diverse population of aggregative organisms, each of which only adheres to clonemates. Because aggregation rate is frequency and density dependent (Figure 6, Supplementary Figures 5, 6), any genotypes that are locally rare will be unable to rapidly form large groups, as they will be capable of interacting with only a small fraction of the population. Strict kin recognition during aggregation therefore undermines the ecological advantage of aggregation. This is even more of a problem if the benefits of aggregation require that a size threshold be met (*e.g.,* enough individuals to form a multicellular fruiting body^52^).

## Discussion

Development is a fundamental aspect of multicellularity, orchestrating the pattern of cellular behaviors that give rise to multicellular phenotype and influencing a lineage’s evolutionary potential. Despite significant theoretical work, the lack of appropriate model systems has limited our ability to directly test the role of developmental mechanism on the subsequent evolution of multicellularity. We circumvent this limitation by engineering aggregative and clonal development from an isogenic unicellular yeast ancestor (Figure 1a).

We grew our yeast under conditions in which selection favored a rapid transition from a unicellular to multicellular stage, the type of environment that is thought to favor aggregative multicellularity^12^. The advantage that aggregative floc yeast showed in monoculture (Figure 2) evaporated once they were competed directly with clonally-developing snowflake yeast (Figure 3), the result of a wholly unexpected social exploitation. Snowflake yeast, which do not produce adhesive Flo1 proteins, embed themselves within large floccy aggregates at a higher frequency than the floc genotype (Figures 3c&d, 5; Supplementary Figures 3, 4). As a result of this social exploitation, snowflake yeast rapidly displace floc (Figure 3b). This result is even more striking in light of prior work in flocculating yeast, where Smukalla *et al* (2008) show that *FLO1* acts as a greenbeard gene, excluding unicellular *FLO1*^−^ competitors from the floc. This is thought to be a consequence of preferential binding between *FLO1*^+^ cells, leading to phase separation^43^. In our case, the ability for *FLO1*^−^ snowflake yeast to co-aggregate with floc appears to arise as a consequence of their branchy structure, allowing them to become entangled within a floc. Our results also provide context for understanding the results of a prior experiment, in which five wild isolates of flocculating yeast were evolved with daily settling selection. Here, snowflake yeast arose *de novo* and largely displaced their floc ancestors in 35/40 replicate populations^40^.

Self / nonself recognition systems play a key role during the evolution of multicellularity, limiting the potential for within-organism genetic conflict^49,50,53^. This may be especially important in lineages that develop aggregatively, as they are more likely to form genetically-diverse multicellular groups. Kin-recognition mechanisms have evolved independently in cellular slime molds^49,53^ and *Myxococcus* bacteria^50,54^, both of which develop via aggregation. We explored the evolution of self-recognition in our system using a mathematical model. We considered our standard ‘permissive floc’, which binds to other permissive floc or snowflake yeast, and ‘choosy floc’, which only attaches to clonemates. While it might seem like choosy floc (which axiomatically cannot experience social conflict) would always be at an advantage, this was not true. Permissive binding increases opportunities for cell-cell adhesion, increasing aggregation speed and group size. Indeed, our experiments show striking support for this hypothesis: floc that formed chimeric aggregates were capable of settling much faster than floc alone (Figure 3a). In our model, choosy floc pay an ecological cost when rare, as it can only bind a small fraction of the cells in the population, forming small groups. This strong positive-frequency dependent selection makes it difficult for choosiness to arise from a population of permissive ancestors, a cost which is compounded if the population is composed of multiple choosy genotypes, each of which is only capable of adhering to clonemates. Consistent with this hypothesis, kin discrimination systems in extant aggregative organisms are quite permissive: wild-collected isolates readily form genetic chimeras^53,55,56^, sometimes (but not always^29,57^) resulting in social cheating^50,58,59^.

Our results highlight a fundamental trade-off faced during aggregative development: selection for rapid group formation often favors permissive binding, but the resulting high within-group genetic diversity lays the foundation for persistent evolutionary conflict. This has important implications for the evolution of multicellular complexity, as the resulting genetic conflict can undermine multicellular adaptation^39^. Indeed, aggregation is relatively uncommon among independently-evolved multicellular lineages^14,60^, and all known examples of independently evolved ‘complex multicellularity’ (*i.e.*, metazoans, land plants, mushroom-forming fungi, brown algae, and red algae^11^) develop clonally. In the context of major evolutionary transitions, aggregation appears to be self-limiting, the evolutionary potential of aggregative lineages constrained by an ecological imperative for effective group formation.

## Methods

### Strain construction

All strains used in this study are listed in Supplementary Table 1. We constructed snowflake and flocculating genotypes from a single clone of the initially unicellular *S. cerevisiae* strain Y55. Snowflake yeast were made as in^61^, but we replaced the *ACE2* ORF with *HYGMX.* Flocculating yeast were made by amplifying the *KAN-GAL1*p*::FLO1* cassette from DNA template from *S. cerevisiae* strain KV210^62,63^ and replacing the *URA3* ORF in our ancestral strain. *ura3Δ::KAN-GAL1*p*::FLO1*/*ura3Δ::KAN-GAL1*p*::FLO1* diploids were obtained by autodiploidization of single spores collected via tetrad dissection onto Yeast Peptone Dextrose plates (YPD; per liter: 20 g dextrose, 20 g peptone, 10 g yeast extract, 15 g agarose) then replica plated onto YPD + 200 mg/L G418. Transformants were confirmed by PCR as well as phenotype when grown in YPGal medium (per liter: 20 g galactose, 20 g peptone, 10 g yeast extract). For microscopy and competition experiments, strains were tagged with green and red fluorophores. To do this, plasmids pFA6a-TEF2Pr-eGFP-ADH1-Primer-NATMX4 and pFA6a-TEF2Pr-dTomato-ADH1-Primer-NATMX4 were amplified and inserted into the *LYS2* locus, and transformants were confirmed via fluorescent microscopy. All transformations were done using the LiAc/SS-DNA/PEG method of transformation^64^.

### Competitive success assay

To determine if snowflake yeast had a competitive advantage over floc yeast, we competed snowflake and floc starting at a range of initial genotypic frequencies (0-100% snowflake in 10% increments) over two days of daily selection for fast settling for 5 min on the bench as in Ratcliff *et al.*, 2012. Snowflake and flocculating yeast were grown up in a mixture of galactose and glucose (YPGal+Dex; per liter; 18 g galactose, 2 g dextrose, 20 peptone, 10 g yeast extract) for 24 h at 30°C, shaking at 250 rpm. This concentration of galactose and glucose was used because it yielded clusters of similar size after 24 h of growth in snowflake and floc yeast (mean floc log(volume) = 12.5, mean snowflake log(volume) = 11.5, *t*(2) = −0.39, *p* = 0.73). Five replicates of 500 µL of each starting genotypic frequency was mixed from overnight cultures and 100 µL of this culture was diluted into 10 mL YPGal+Dex for the competition experiment. The remaining 400 µL was used to measure the initial count of snowflake and floc yeast. To do this, EDTA (50 mM, pH 7) was used to deflocculate cells to run through a CyFlow® Cube8 flow cytometer where two distinct peaks corresponding to unicellular floc cells and snowflake cultures could be counted. Counts of unicellular floc and snowflake yeast were obtained for time 0 and after three days of competition. The competitive success of snowflake yeast was calculated as the ratio of snowflake to floc yeast after competition relative to before competition using the equation (1):

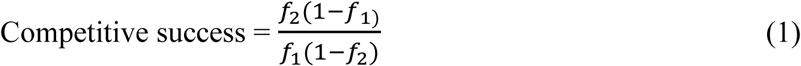

where *f_1_* is the frequency of snowflake yeast before competition and *f_2_* is the frequency of snowflake yeast after competition^37^. This fitness measure is simple and general (*i.e.*, it doesn’t assume any underlying model of population dynamics, like exponential growth), and accommodates different starting frequencies.

### Measuring settling rate

Unlike snowflake yeast, floc yeast form groups as they are settling, so we needed to measure the properties of flocs during the process of settling directly. To do this, we developed a novel, robust, high-throughput method of measuring the settling speed of yeast populations. Various methods to measure aggregation and settling in yeast exist^24,65–68^, but most of them introduce experimental variables that limit their relevance to our system^65^, and no method is considered standard in yeast research in general^65,68,69^. Importantly, most of them lack the temporal resolution needed (seconds) to capture the fast-settling profiles of some of our strains. In our method, we placed the yeast in back-illuminated cuvettes, and used high-speed high-resolution video acquisition (24 fps, 3840 × 2160 pixels, Sony a7R II, 90 mm macro lens) to capture changes in pixel densities over the settling time (Figure 1b). Our method relies in the fact that settling and flocculation produce optically denser regions, relative to the initial density distribution (Supplementary Movie 1), thus allowing us to measure the rate of this density changes. We pre-processed our raw density data with a Savitzky-Golay smoothing function in order to preserve the signal over the noise without sensibly changing the shape of the dynamics (Supplementary Figure 10). We then calculated a characteristic settling rate, as the maximum slope in the density dynamics. We validated our method by quantifying the percentage of biomass settled at 5 min in floc and snowflake cultures, showing that, as expected, a higher settling rate indicate a higher proportion of settled cells (Supplementary Figure 11).

### Competitive success during growth and settling

There are two important life history traits in our experimental system: growth rate and settling rate^45,47^. We measured the competitive success of snowflake yeast during both stages. Snowflake and floc yeast were grown separately for 24 h in YPGal+Dex. As above, five replicates of 500 µL of various starting genotypic frequencies (10-90% snowflake in 20% increments) were mixed from overnight cultures and 100 µL was used to dilute into fresh YPGal+Dex and the remaining 400 µL was used to calculate initial snowflake and unicellular floc counts as described above. To measure snowflake competitiveness during growth, 500 µL of each culture was deflocculated using EDTA and snowflake and floc counts were measured on the flow cytometer after 24 h or growth at 30°C, shaking at 250 rpm. To measure competitive success over one round of settling selection, 2 mL of each snowflake/floc co-culture was aliquoted into 2 mL microcentrifuge tubes. 500 µL was then aliquoted into 1 mL microcentrifuge tubes and deflocculated to obtain pre-selection snowflake and floc concentrations as described above. The remaining 1.5 mL was allowed to settle on the bench for 5 min, after which the top 1.4 mL was discarded. The remaining 100 µL was deflocculated and post-selection snowflake and floc counts were obtained via flow cytometry.

### Examining the composition of aggregates

The composition of snowflake and floc yeast within large chimeras was measured by fluorescent microscopy, using a Nikon Eclipse Ti inverted microscope with a computer-controlled Prior stage. Specifically, snowflake and floc-GFP were grown for 24 h in YPGal+Dex. Four replicates of snowflake and floc co-cultures with differing amounts of starting snowflake (20% or 80%, respectively) were inoculated into fresh medium and grown for another 24 h. 10 µL of this culture was placed between a slide and a 25 × 25 mm coverslip and the whole coverslip was imaged by combining 150 separate images at 100 × magnification yielding a 42456 × 42100 pixel (1.78 billion pixels; 1.23 × 1.22 cm) composite image. The percentage of biomass in different cluster size classes belonging to either snowflake or floc yeast was calculated using a custom Python script. “Large flocs” were considered to be anything larger than the largest snowflake clusters (Figure 5c).

### Mathematical modeling

We consider a settling competition between snowflake clusters and flocculating cells. If flocculation, settling, and reproduction all occur together we might expect a complicated set of dynamics resulting from the interplay between these processes. We simplify our analyses by focusing strictly on aggregation. We assume that aggregation and settling happen after the primary growth phase and occur faster than reproduction such that the populations of cells are large as a result of several generations of reproduction in media. Furthermore, we consider aggregation and settling as two separate processes. Thus, we assume that there is some time in which cells aggregate and afterwards the groups are exposed to settling selection. This assumption allows us to focus on modeling the dynamics of aggregation and circumvent explicit spatial models that would be required to consider the dynamic interactions between aggregation and settling via centrifugation. We model the dynamics of aggregation using a system of differential equations, where a snowflake cluster composed of *i* cells is denoted as *S_i_*, a floc of *i* choosy cells is *C_i_*,and a floc of *i* permissive cells is *P_i_*. (equations 2-5):

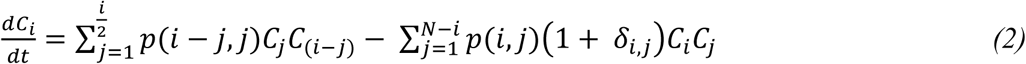

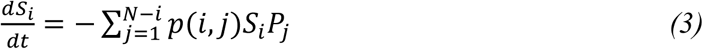

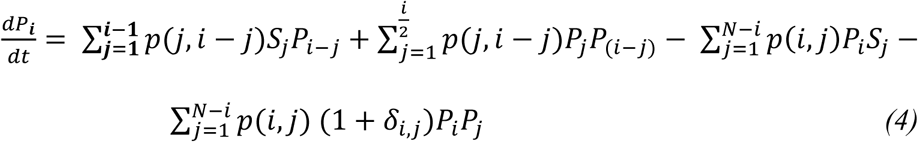

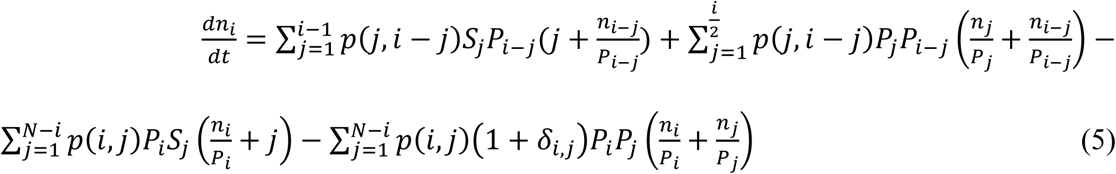

Choosy floc clusters *C_i_* can bind to former larger flocs but we assume each cluster reaches a maximum size *N* (*N*=1000 in our computations). We also ignore group fragmentation. Thus: 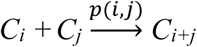, where *i* + *j* ≤ *N* and *p(i,j)* is the probability of a successful binding that depends on the size of two flocs. Specifically:

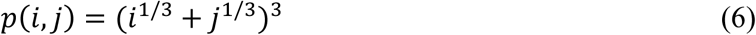

This function depends on many factors including the geometry of the two clusters, the probability of collision, the probability of a collision resulting in binding, etc. We assume that it is a simple function of the radii of the two clusters: *p(i,j)* = *(r_i_* + *r_j_)^3^* where *r_i_* and *r_j_* are the radii of *C_i_* and *C_j_* and the radii can be approximated by considering the clusters as spheres. Thus, if the volume of a single cell is 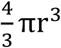, then the volume of *C_i_* is 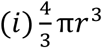 which makes the radius of *C_i_* equal to *i*^1/3^*r*. We consider *r* = 1 to simply the calculations. The *δ_i,j_* term accounts for the extra loss if two identically-sized flocs interact, i.e. if two *C_i_* bind then the loss is double that of a *C_i_* binding a *C_j_* where *i ≠ j*.

We use *P_i_* to denote permissive flocs. Since *P* cells can bind to either its own cells or snowflake cells, a *P_i_* cluster may be composed of k floc cells and *i* – *k* snowflake cells for any *k* ≥ 1. We assume that there are a large number of clusters and cells and track the number of snowflake cells in *P_i_* clusters for each size *i*, which we denote *n_i_*. This assumption corresponds to treating the aggregative mixture as a classic tank mixing problem.

In all competitions except for Supplementary Figures 8&9, we assume an initial inoculum of 1000 concentration units that is split between *C*, *S*, and *P*. The initial distribution of *S_i_* is fit to a lognormal distribution that matches empirical data (Supplementary Figure 5). This distribution only changes in the presence of permissive floc. The winner of the settling competition is determined by solving equations (2-5) for some time *t* and selecting the largest 10% of the population, using group size as a proxy for settling speed. This is analogous to our experimental system, where 10% of fastest-settling yeast biomass gets passaged to the next tube following settling selection. For *C* cells, as time increases, more of the distribution is represented in the largest fractions (≈ *i* = *N*; Supplementary Figure 5). Thus, the amount of *C* cells in the top 10% of possible clusters size increases with time, but levels out for longer *t* (Supplementary Figure 5).

The mathematical model captures a single round of aggregation and selection without regard to how populations grow in between selective events. In cases where we consider multiple rounds of aggregation and selection (Supplementary Figure 8), we do not use any explicit models of population growth. Rather, we multiply the final proportions of cells after selection by the inoculum size and use that as the input to the next iteration of aggregation and selection. This bypasses the possibility that different population growth dynamics might alter the proportions of cell types. In addition, we also assume that the *P* and *S* cells dissociate from their mixed groups and begin the next aggregation and selection phase as separate entities.

## Author Contributions

J.P., W.C.R. and P.M.Z designed the experiments. J.P. conducted the experiments. J.P. and P.M.Z. designed the settling rate analysis pipeline. All authors contributed to data analysis and interpretation. E.L. performed the modeling. P.Y. performed the spatial analysis of snowflake yeast within chimeric aggregates. J.P. and W.C.R. wrote the first draft of the paper, all authors contributed to revision. This work was supported by NASA Exobiology grant no. NNX15AR33G to W.C.R. and E.L., NSF grant no. IOS-1656549 to W.C.R. and P.J.Y., NSF grant no. IOS-1656849 to E.L., and a Packard Foundation Fellowship for Science and Engineering to W.C.R.

## Acknowledgements

We would like to thank Kevin Verstrepen for providing us with strain *S. cerevisiae* KV210, which contained the *GALp::FLO1* construct, and Daniel Weissman for insightful feedback on the manuscript.

