## Supplementary Information for "Ecological advantages and evolutionary limitations of aggregative multicellular development"

### Supplementary Methods

#### Direct measurement of settled biomass

To validate our imaging-based approach for quantifying settling speed, we measured biomass settling for floc and snowflake yeast directly. Five replicates of floc or snowflake yeast were grown up in 10 mL YPGal+Dex for 24 h at 30°C, shaking at 250 rpm. 1.5 mL of stationary-phase cultures were placed in a pre-weighed 2 mL microcentrifuge tube. Yeast were then allowed to settled for 5 min at 1 g, after which the top 1.4 mL was transferred to a different pre-weighed tube. Yeast were pelleted and double-washed with deionized water, then the excess water was removed and the pellet was airdried 50°C for two days. Settling rate was determined as the percentage of total biomass in the bottom 100µL pellet.

#### Measuring the ratio of flocculating and non-flocculating unicells in flocs

We measured the ratio of flocculating (*FLO1*) and non-flocculating (*flo1*) cells in flocs to determine of *flo1* cells are preferentially excluded. *FLO1*-GFP and *flo1* cells were grown separately for 24 h in YPGal+Dex. Three replicates of *FLO1*-GFP and *flo1* cocultures with a starting ration of 90:10 *FLO1*-GFP : *flo1* or 50:50 *FLO1*-GFP : *flo1* were inoculated into fresh medium and grown for another 24 hours. Flocs were separated from planktonic cells and the flocs were deflocculated with EDTA. The number of cells of each type in the floc and planktonic populations were measured via flow cytometry.

**Supplementary Table 1. Strains used in this study.**

| **Strain** | **Relevant Genotype** | **Reference** |
|---|---|---|
| Snowflake | Δ*ace2*::HYGMX | This study |
| Floc | Δ*ura3::KAN-GAL1*p::*FLO1* | This study |
| Snowflake-GFP | Δ*lys2*::*TEF2*p-yeGFP | This study |
| Snowflake-RFP | Δ*lys2*::*TEF2*p-dTomato | This study |
| Floc-GFP | Δ*lys2*::*TEF2*p-yeGFP | This study |
| Floc-RFP | Δ*lys2*::*TEF2*p-dTomato | This study |

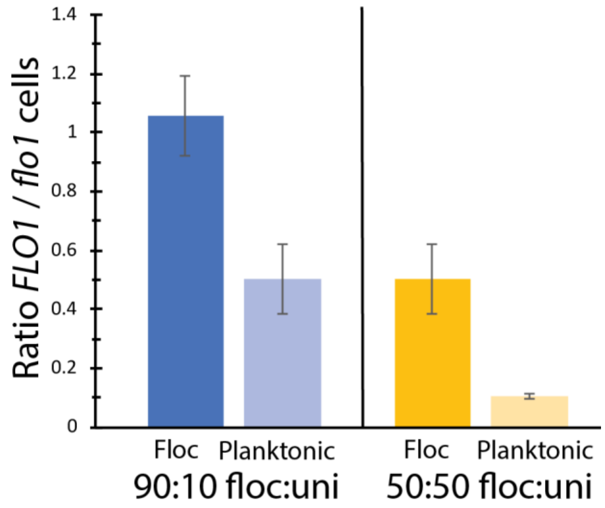


**Supplementary Figure 1. Non-flocculating unicells are largely excluded from flocs.** Shown are the ratio of flocculating (*FLO1*) to non-flocculating (*flo1*) cells in the flocculating and plaktonic subpopulations of co-cultures. These populations were initially inocluated at a ratio of 90:10 *FLO1:flo1* or 50:50 *FLO1*:*flo1* cells. *flo1* cells are preferentially excluded from flocs. Error bars are standard deviations of three biological replicates.

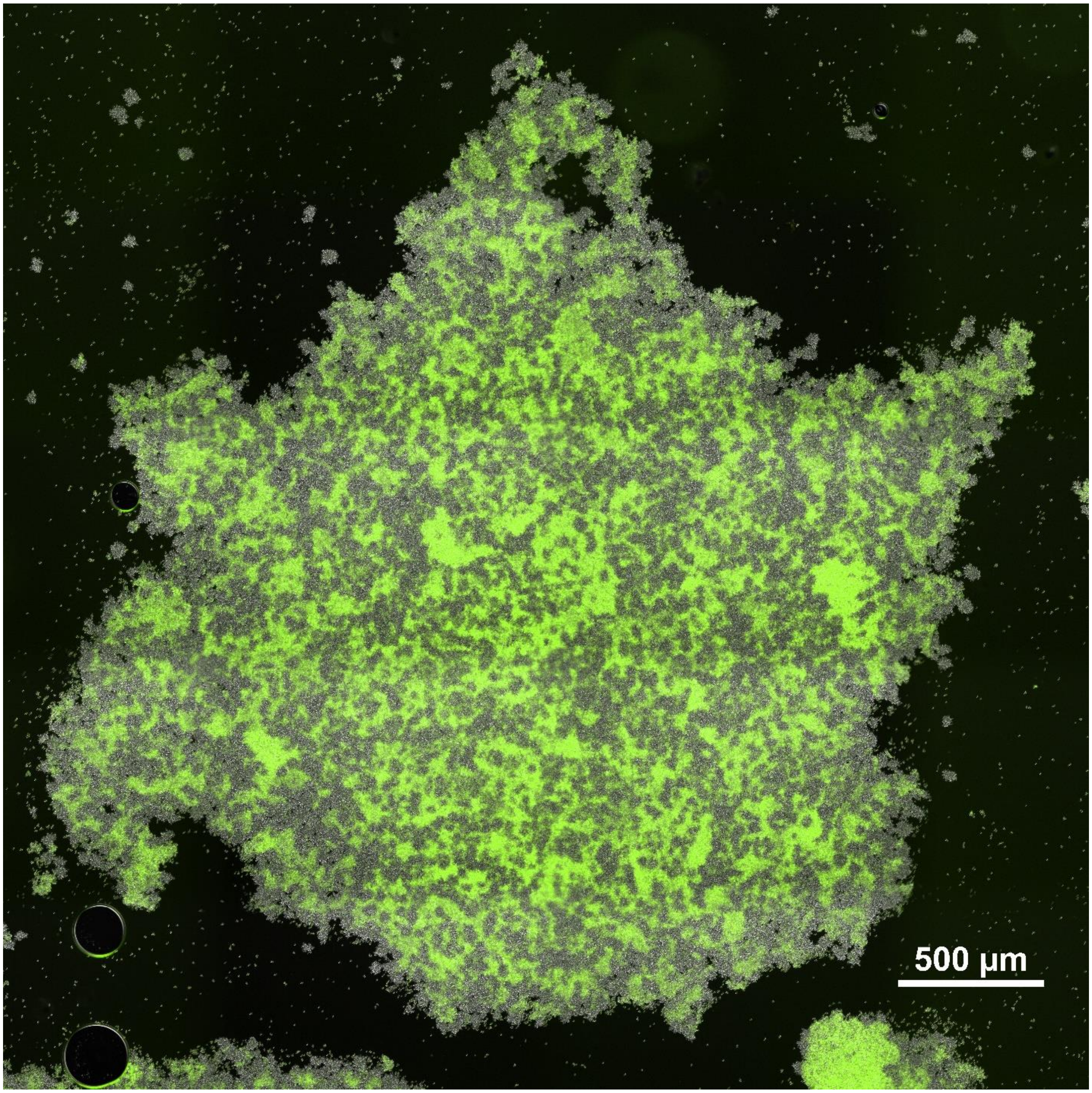


**Supplementary Figure 2**. Large chimeric aggregate from a population that is 70% floc, 30% snowflake yeast.

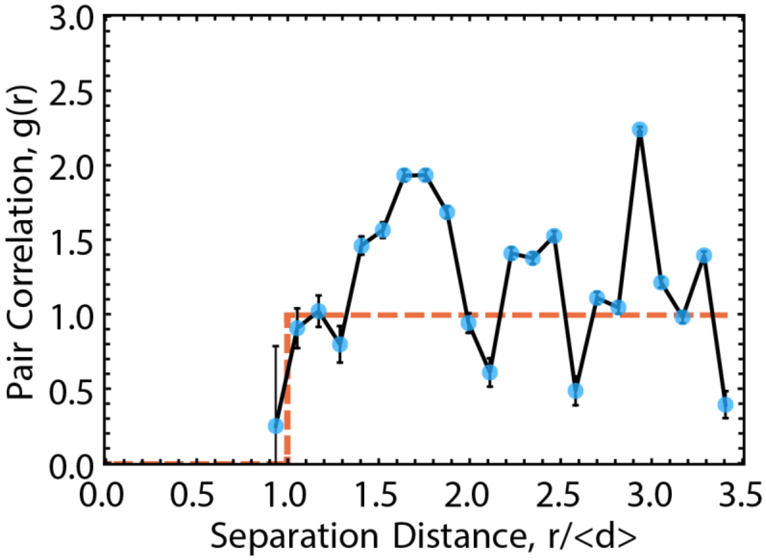


**Supplementary Figure 3. Pair correlation function measurement.** The pair correlation function, *g(r)*, is measured from snowflake yeast cluster positions within aggregates, taken from micrographs. The black line represents the pair correlation function measured in a sample that is 10% snowflake yeast with standard error bars. The red line represents the pair correlation function for a random distribution of clusters.

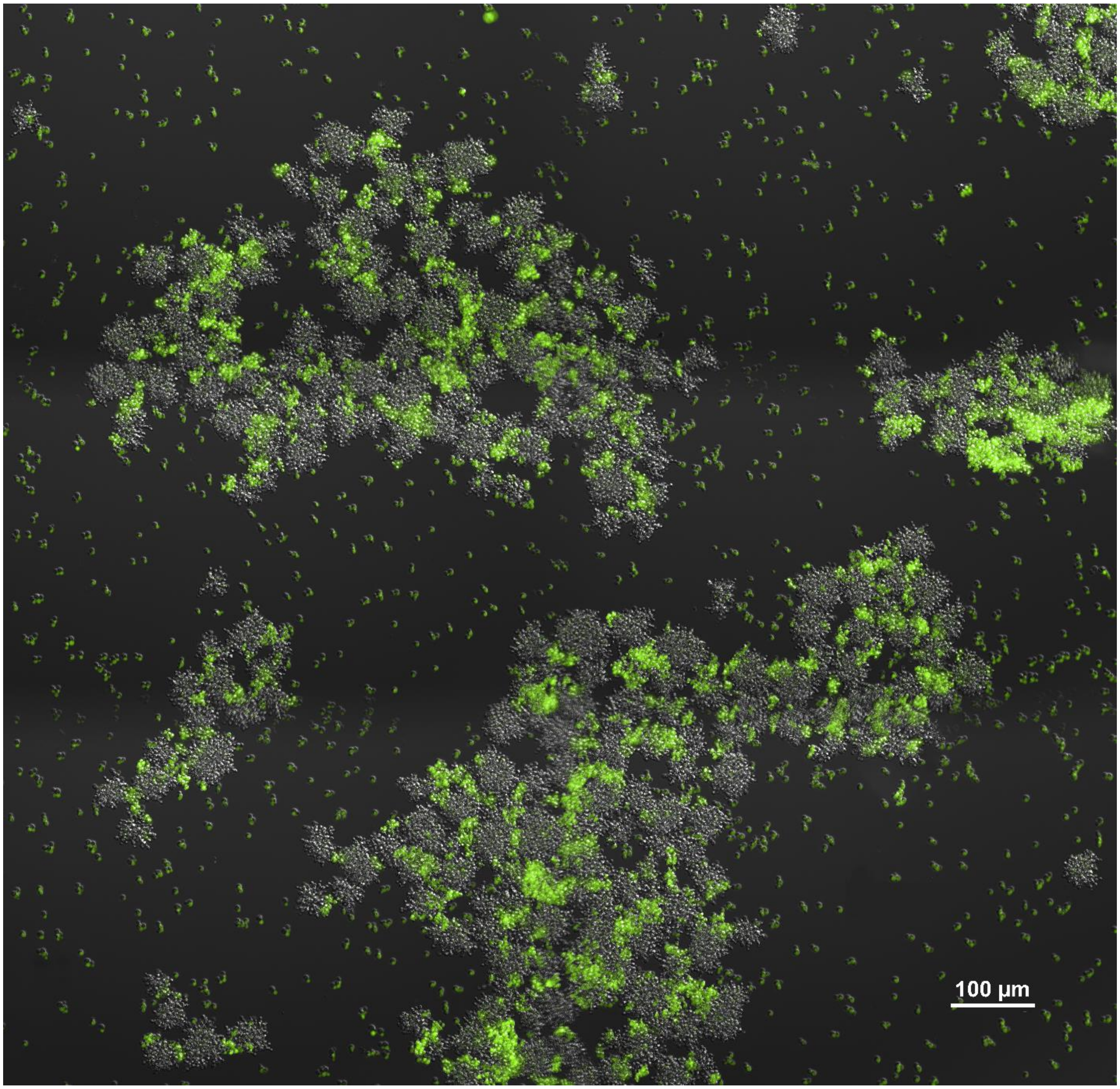


**Supplementary Figure 4. Floc yeast act as an adhesive, allowing snowflake yeast to form aggregative chimeric aggregates.** This is an image from a population with 30% floc, 70% snowflake yeast. All snowflake yeast within chimeric aggregates adhere via small patches of floc cells (labeled with GFP).

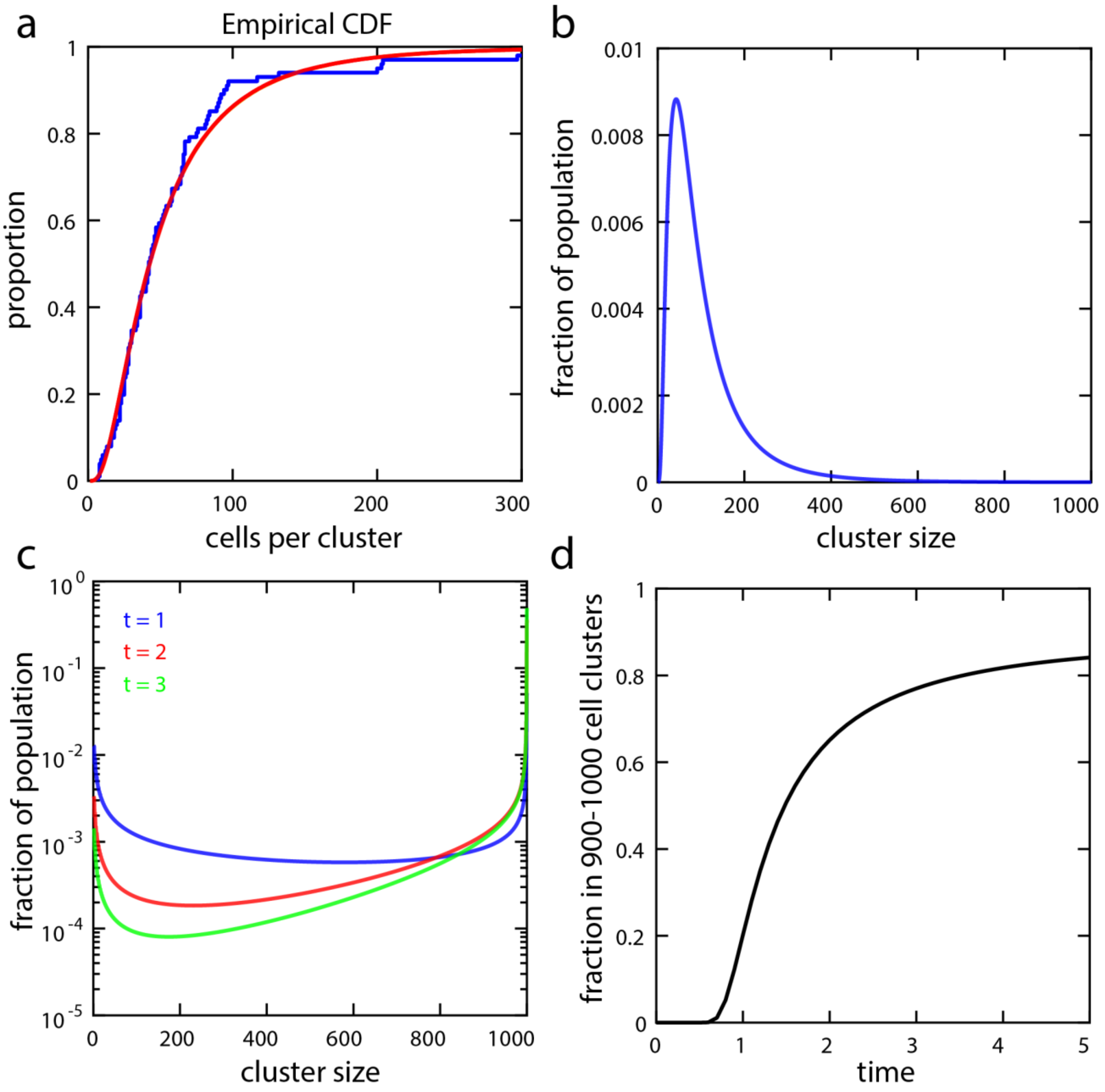


**Supplementary Figure 5. Distribution of $S_i$ and $P_i$ as a function of $i$.** a) The blue curve is the cumulative density function of the distribution of snowflake cluster sizes using empirical data and the red curve is the fit to a lognormal distribution. b) The distribution of snowflake cluster size $S_i$ used in model simulations, which does not change over time. c) Distribution of permissive floc cluster sizes $P_i$ for three different times. As the amount of time for aggregation increases, a greater proportion of the distribution is represented in the largest cluster size fractions. d) The fraction of $P_i$ in the top 10% of possible clusters sizes increases with aggregation time, leveling out for larger $t$ as $P_i$ reaches the maximum cluster size.

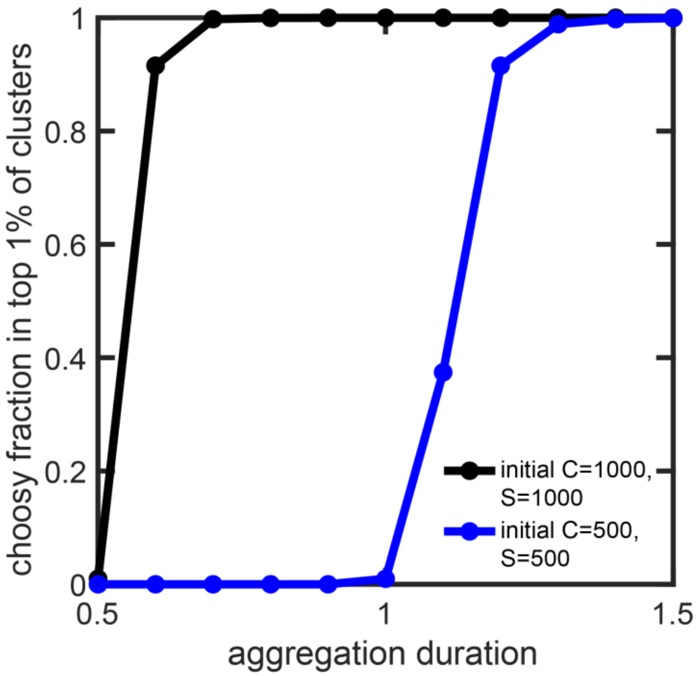


**Supplementary Figure 6. Effect of population size on the aggregative fitness of choosy floc, *C*.** Choosy floc and snowflake do not co-aggregate during competition. We can thus examine the effect of overall population size on *C*'s aggregation by varying population size (increasing the density of *C* and probability they will interact and aggregate), and aggregation duration. Higher densities and longer aggregation durations favor choosy floc.

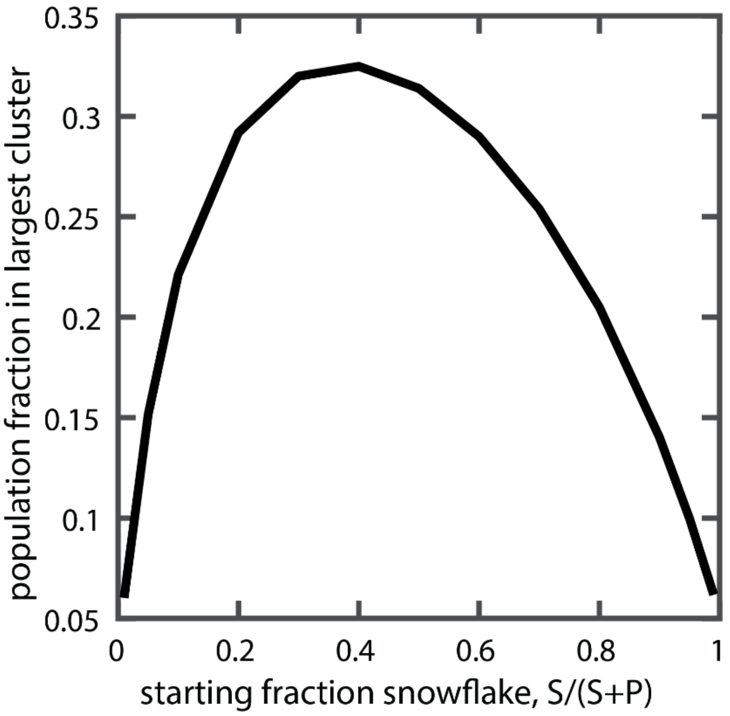


**Supplementary Figure 7. Populations of snowflake and permissive floc yeast form large, fast-settling aggregates at intermediate frequencies (peaking at 40%).** This is similar to experimental data showing a peak settling speed between 30 and 40% *S* (Figure 3a).

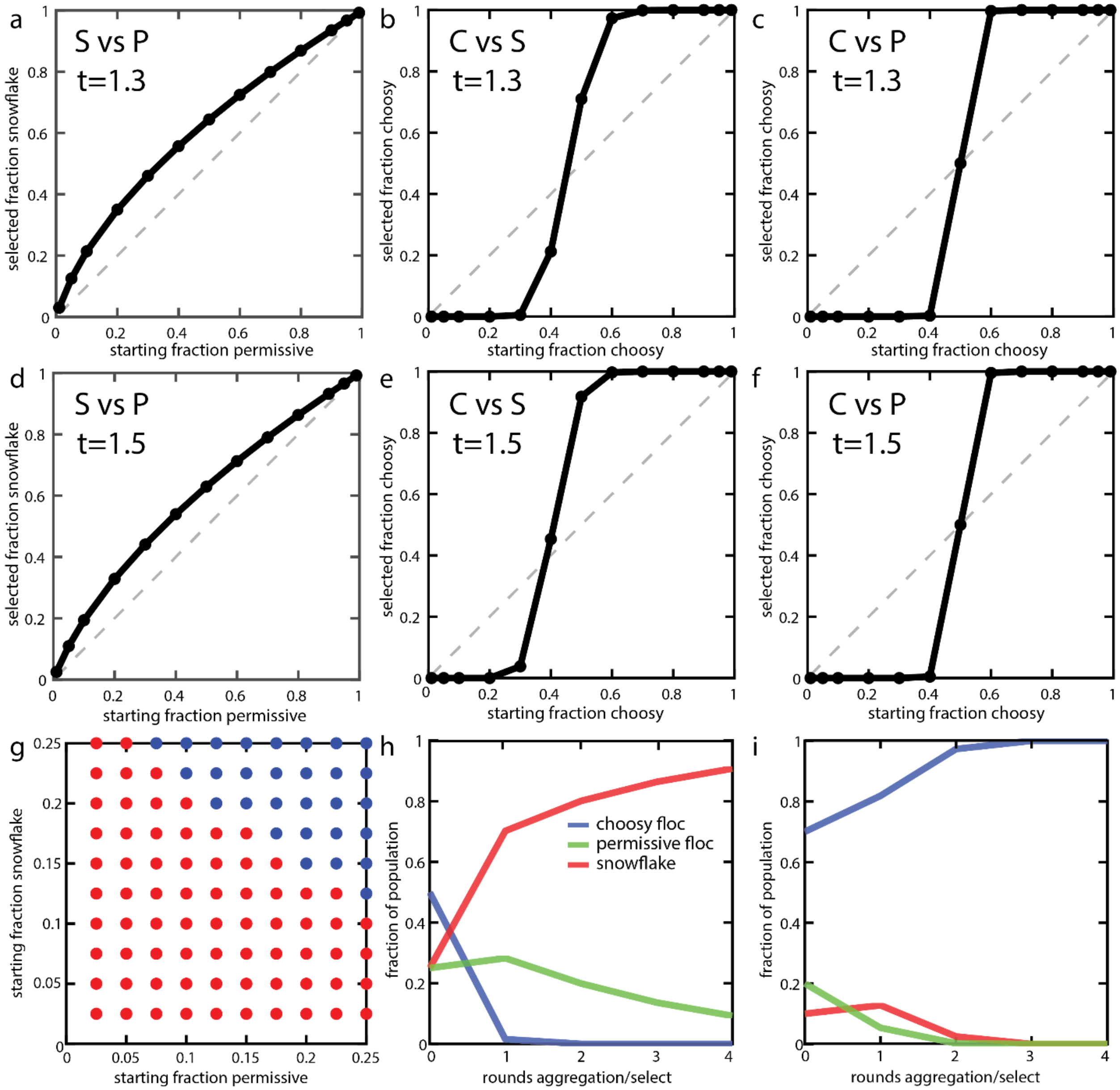


**Supplementary Figure 8. Effect of aggregation time on model dynamics.** We examine the dynamics of all three pairwise competitions between *S*, *P* and *C* as a function of aggregation time; *t*=1.3 (a-c) and *t*=1.5 (d-f). In general, longer aggregation times favor choosy floc, giving it a greater opportunity to form large clonal aggregates. a-f can be compared to Figure 6a-d, where *t*=1. g) Invasion phase diagram for the three-way competition, with *t*=1.5; when red, *C* increases in frequncy. Note that at this time duration, *C* can displace *P*+*S* from a lower starting frequency than when *t*=1 (Figure 6d). h & i) Competition across multiple rounds of settling selection under conditions favoring *C*, namely, long aggregation (*t*=1.5) and strong selection (biomass from the largest groups composing 1% of population's biomass survive settling selection). *C* are unable to displace *P* and *S* when starting at 50% frequency (h), but are when starting at 70% (i). Note also how over the first timestep in (h), snowflake yeast and permissive floc both increase in frequency, then snowflake yeast subsequently parasitizes the floc.

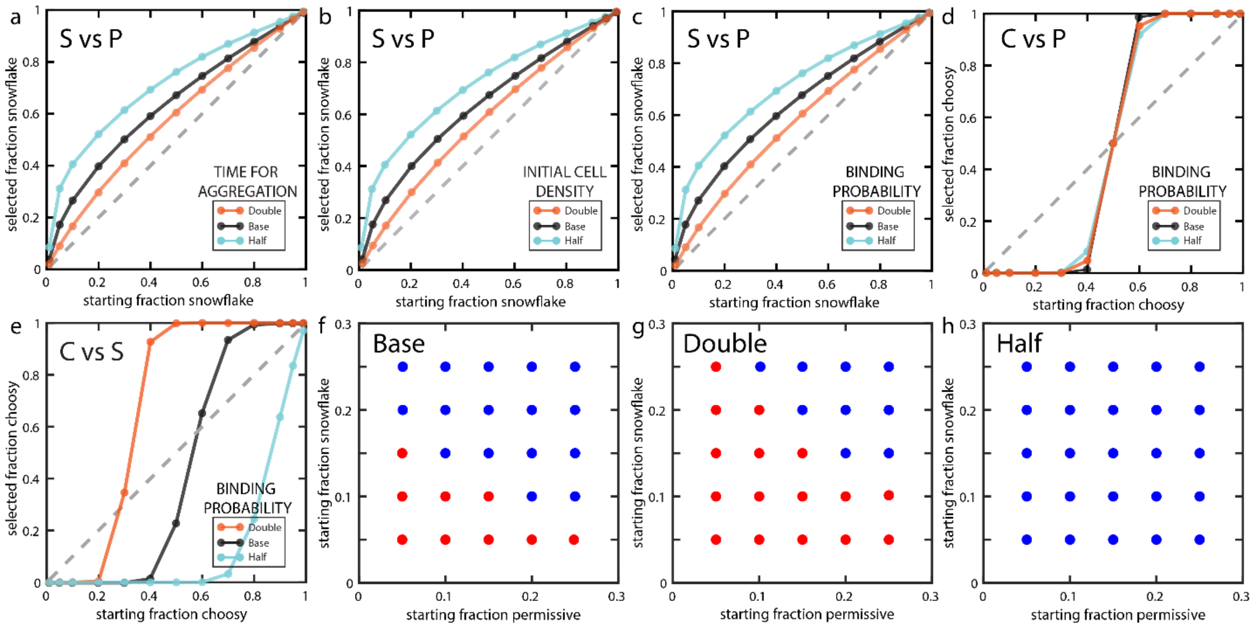


**Supplementary Figure 9. Parameter sensitivity analysis.** We analyzed the dependence of our model on three key parameters (aggregation time, initial cell density, and binding probability) by either doubling or halving each parameter relative to the base calculation (Figure 6). a-c) Increasing aggregation time, initial cell density, or binding probability gives an advantage to *P* cells over *S*, but *S* still increases in relative frequency at all initial starting *S* fractions. Additionally, when each of these parameters are either doubled or halved, the result is the same. Thus, our model has the same sensitivity to each of these parameters. d) *C* vs *P* yields the same result regardless of how the parameters change because *C* and *P* flocs are identical in the absence of *S*. e) *S* cannot be invaded if the binding probability (or aggregation time or initial cell density) is halved, but only wins when it starts at more than 75% of the population when doubled. f-h) Invasion diagram for three-way competition when the binding probability (or aggregation time or initial cell density) is halved or doubled. When red, *C* increases in frequency. If parameters are doubled (g), favoring flocculation types, *S*+*P* still win at some frequencies, but *S*+*P* always win when parameters are halved (h).

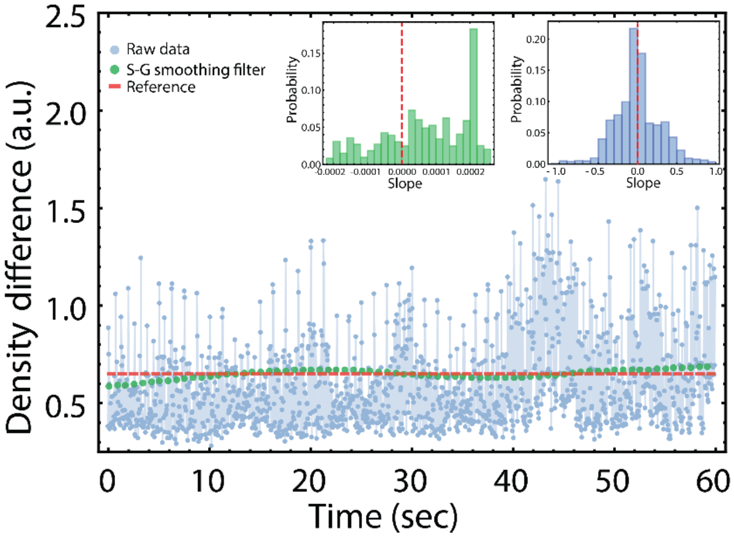


**Supplementary Figure 10. Processing of raw density data.** Raw data (dotted blue line) from our density measurements have small fluctuations (note the range on the y-axis, ~200 times smaller than that of Figure 1B) that nevertheless can be problematic to estimate a characteristic 'maximum slope' of the overall dynamics. We show here a negative control, spent media, where we expect no significant changes in density over time, corresponding to a characteristic slope of zero (red dashed line, intersect in Y at 0.652 corresponds to the average of the dynamics). Using a Savitzky-Golay smoothing function with a window size of 415 (various window sizes give similar results), we were able to recover a distribution of slopes very close (within 2 x $10^{-4}$) to what was expected (green histogram), compared to the raw data (blue histogram). This process yielded a high signal-to-noise ratio (28.04 compared to 2.19 of the raw data - defined here as $\mu/\sigma$, the reciprocal of the coefficient of variation). Therefore we applied the same pre-processing to all our raw density data. This process was implemented with MATLAB built-in functions (MATLAB, MathWorks).

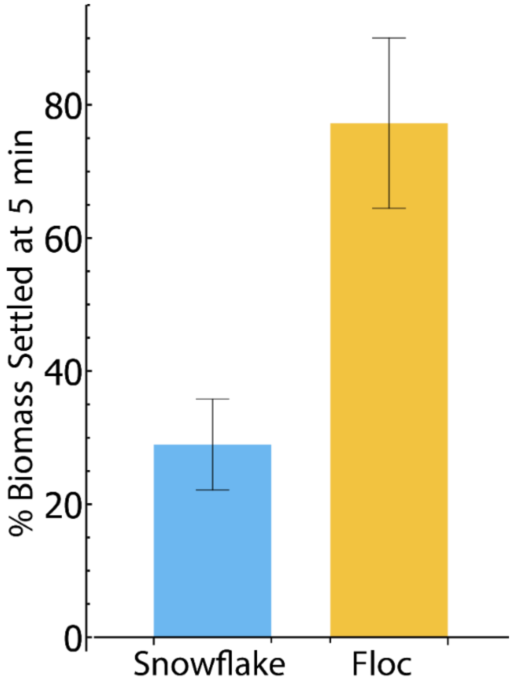


**Supplementary Figure 11. Biomass measurement of settling survival.** We verified that floc yesat settle more rapidly than snowflakes, as determined by our biomass-displacement method (*i.e.* Figures 1b, 2a, Supplementary Movie 1), by directly measuring the amount of yeast biomass to survive the 5 minute settling rate challenge. Floc settles roughly 2.65 times better over 5 minutes of settling than snowflake yeast (77% vs 29% settled, respectively; $t_5 = 7.44$, $p$=0.0003, two-tailed t-test). Error bars represent standard deviation of five biological replicates.

**Supplementary Movie Figures**

**Supplementary Movie 1.** Individual pixel intensities, termed here Focal Densities (in the [0, 1] interval), were measured over 5 minutes of settling, at 24 frames per second (see Methods section for details). At all frames and for all focal densities, the absolute difference, relative to the first frame, was quantified. The total of these differences (for the entire cuvette at each frame) was expressed as the density change. We show two representative settling dynamics, corresponding to one of the fastest (80:20 Floc:Snowflake) and one of the slowest (10:90 Floc:Snowflake) settling rates. We show a cell-free negative control with no expected density changes (spent media), showing that there is no overall change in the density. The raw data is shown as shadowed lines in the graph. To correct for the noise in the raw data, we applied a Savitzky-Golay filter (see Supplementary Figure 10 for details), corresponding to the dashed lines of the graph.

**Supplementary Movie 2.** 5 minutes of settling in a monoculture of snowflake yeast (left) and floc yeast (right).
